# *MCH4* is a multicopy suppressor of glycine toxicity in *Saccharomyces cerevisiae*

**DOI:** 10.1101/653444

**Authors:** Artem V. Melnykov, Elliot L. Elson

## Abstract

*Saccharomyces cerevisiae* can either import amino acids from the surrounding or synthesize inside the cell, and both processes are tightly regulated. Disruption of such regulation can result in amino acid toxicity to the cell through mechanisms that are poorly understood. In this study we make use of a mutant strain with deregulated general amino acid permease gene whose growth is inhibited by low concentrations of several amino acids. We carry out multicopy suppression screen with several toxic amino acids and identify *MCH4* as a gene that suppresses inhibitory effects of glycine. We find that expression of *MCH4* is regulated by osmotic shock but not other kinds of stress. These findings are discussed in the context of possible mechanisms of amino acid toxicity.

## Introduction

Amino acids are important metabolites for every cell. In yeast, the general control system (GCN) responds to deprivation of any amino acid by activating transcription of biosynthesis genes. At the same time, the nitrogen discrimination pathway regulates acquisition of extracellular amino acids to be used as a source of nitrogen. In addition, extracellular amino acids activate expression of specific amino acid transporters via the SPS system which serves the needs of protein synthesis. These systems provide the basis for regulation of metabolism under conditions of amino acid limitation (Zaman et al., 2008).

There is also an opposing tendency which prevents hyperaccumulation of amino acids or imbalance of their concentrations in the cell. Biosynthesis pathways are often inhibited by their end products which is known as feedback inhibition; tryptophan (Miozzari et al., 1978; Braus, 1991) and leucine (Kohlhaw, 2003) biosynthesis are two examples but there are many others (Ljungdahl and Daignan-Fornier, 2012). Additionally, transporters of extracellular amino acids can be downregulated by their substrates which limits influx of amino acids into the cell. The most extensively characterized case is post-translational regulation of the general amino acid permease (Cain and Kaiser, 2011) but this mechanism is very likely widespread.

Yeast can also regulate the intracellular amino acid pool by shuttling amino acids between the cytoplasm and the vacuole since this organelle serves as a repository of amino acids and mediates protein turnover in autophagy. Vacuolar amino acid transport has been studied less than transport across the plasma membrane but several amino acid transporters and exchangers have been identified (Russnak et al., 2001). Finally, yeast are able to export amino acids to the extracellular medium under conditions of overproduction or uncontrolled uptake (Casalone et al., 1997; Feller et al., 1999; Hess et al., 2006; Melnykov, 2016). Presence of mechanisms limiting accumulation of amino acids suggests that while these metabolites are necessary for proper functioning of the cell, high or unbalanced concentrations can have a deleterious effect.

Amino acid toxicity has been reported in budding yeast under conditions of unregulated uptake through the general amino acid permease (Gap1p) (Risinger et al., 2006) and several other broad specificity transporters (Ruiz et al., 2017). Regulation of *GAP1* is understood in greatest detail; it is regulated transcriptionally by the quality of the nitrogen source and posttranslationally by intracellular amino acid concentrations. These modes of regulation can be disrupted by overexpressing the protein from a constituitive (*P*_ADH1_) promoter and, in addition, mutating lysines in positions 9 and 16 to arginines. When grown on a synthetic medium without amino acids such unregulated mutant strain *P*_ADH1_-Gap1p^K9R,K16R^ (from now on referred to as *GAP1*^*UR*^) does not have an obvious growth phenotype. However, its growth is inhibited by addition of 3 mM concentrations of individual amino acids, most notably those that yeast cannot catabolize (lysine, histidine, and cysteine) or catabolize poorly (glycine), but also isoleucine and glutamate. Generally, as Gap1p is a transporter of broad specificity, all natural amino acids except alanine inhibit growth of the *GAP1*^*UR*^ strain to some degree (Risinger et al., 2006). Similarly, overexpression of *AGP1, BAP2, CAN1, DIP5, GNP1, LYP1, PUT4*, or *TAT2* from a multicopy plasmid resulted in amino acid sensitivity (Ruiz et al., 2017). In each case, several amino acids were found to inhibit growth, and the spectrum of inhibitory amino acids was specific to the transporter reflecting known and newly discovered transport specificities. While the authors of these studies did not decipher the mechanism of amino acid toxicity, they ruled out disruption of pH gradient accompanying transport as a plausible explanation. They also showed that it is the amino acid imbalance in the cell and not necessarily the high intracellular concentration that is toxic since an equimolar mixture of four most toxic amino acids did not inhibit cell growth (Risinger et al., 2006).

We decided to exploit these unique observations and identify multicopy suppressors of amino acid toxicity hoping to uncover the underlying molecular mechanism. We identified *MCH4*, a gene with no previously assigned function, as one that confers resistance to glycine in the *GAP1*^*UR*^ background. Below, we provide a detailed account of this finding as well as results of several additional experiments aimed at understanding the mechanism of amino acid toxicity.

## Materials and Methods

### Strains and media

All yeast strains used in this study were constructed in the S288c background from yJF176 (Table 1). Integrative replacement was carried out based on recombination driven by short flanking homology following a published protocol (Becker, D. M. and Lundblad, Victoria, 1993). The growth medium for propagation of yeast strains was YPD (1% yeast extract, 2% peptone, 2% glucose). For assays described in Figures 1 and 2, AmmSC medium was used (0.67 g/l yeast nitrogen base without amino acids, 0.5% ammonium sulfate, 2% glucose). When indicated, ammonium sulfate was replaced with allantoin (0.1%), L-proline (10 mM) or glycine (1.5%), and the resulting media are referred to as AllSC, ProSC, and GlySC, respectively. All medium components and amino acids were purchased from Sigma.

**Table 1.**
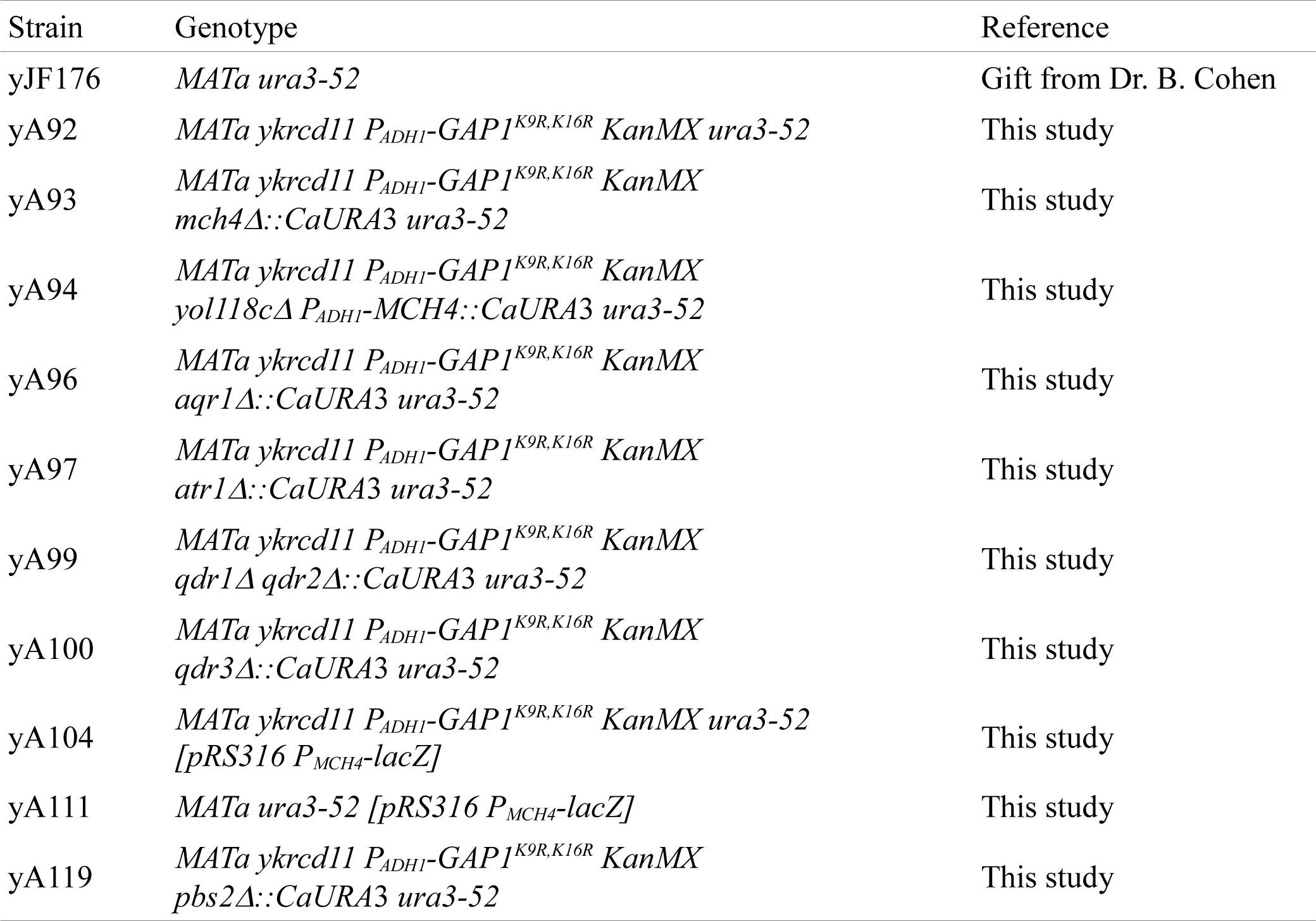
Yeast strains used in this study

**Figure 1.**
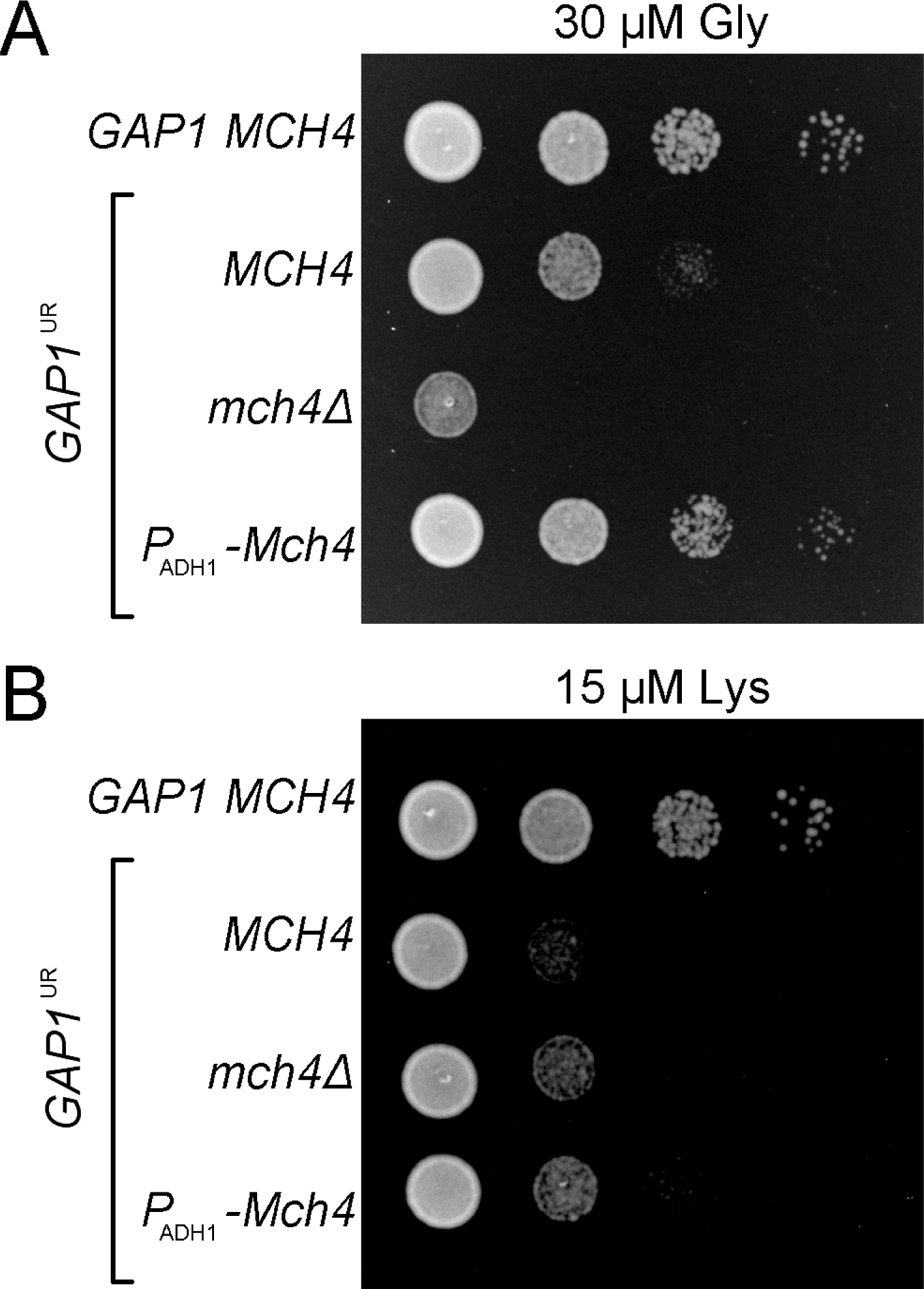
Effect of *MCH4* on toxicity of low concentrations of amino acids (A, 30 μM glycine; B, 15 μM lysine). For each panel, the rows of dilutions from top to bottom are the wild type strain (yJF176) followed by strains sensitive to amino acids (*GAP1*^*UR*^) and varying in *MCH4* locus: wild type *MCH4* (yA92), deleted (yA93), and overexpressed from the *ADH1* promoter (yA94). The cells were grown on complete synthetic medium (AmmSC) supplemented with low concentrations of either glycine or lysine.

**Figure 2.**
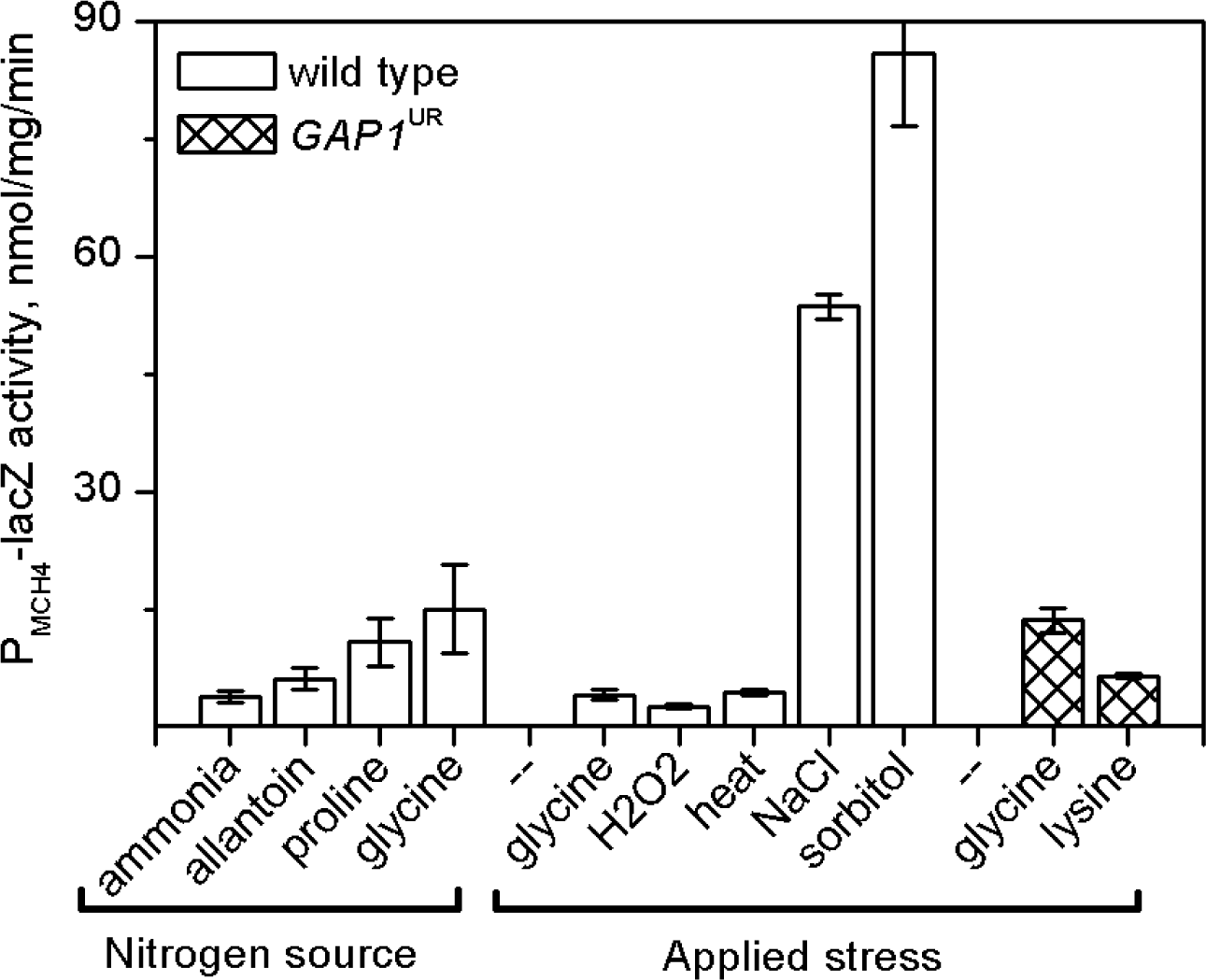
Activity of the *MCH4* promoter as a function of supplied nitrogen source or applied stress for wild type (yAM111) and *GAP1*^*UR*^ (yAM104) strains expressing β-galactosidase from *P*_*MCH4*_ on a low copy plasmid. To study the effect of nitrogen source, the wild type strain was grown on AmmSC, AllSC, ProSC, and GlySC. To study the effect of stress, yeast were grown on AmmSC, and the following stressing factors were applied for 90 minutes: glycine (5 mM for wild type, 500 μM for *GAP1*^*UR*^), lysine (500 μM), H_2_O_2_ (400 μM), NaCl (1 M), sorbitol (1.5 M), and 42 °C incubation. The error bars represent standard deviation of the mean resulting from three independent measurements.

For plate growth assays, yeast were grown to OD_600_ = 1.0 on AmmSC, concentrated by centrifugation, washed once with water, and spotted on the plate as 5 μl of cell suspension of OD_600_ = 0.3, 0.03, 0.003, and 0.0003. All plates were photographed after 42 hour incubation at 30 °C. All spotting assays were repeated at least twice, and gave rise to reproducible growth patterns.

### β-galactosidase activity measurements

A batch culture of the strain of interest was grown to OD_600_ = 0.5 in AmmSC (unless otherwise indicated) at which point glycine, lysine, or H_2_O_2_ was added. Alternatively, the culture was grown to OD_600_ = 1.0 and mixed with an equal volume of the same medium containing 2 M NaCl or 3 M sorbitol. The cells were incubated for 90 minutes after the addition of the stressing factor. For studying the effect of the heat shock, the yeast batch culture was combined with an equal volume of the medium prewarmed to 49 °C, and incubation was continued for 90 minutes at 42 °C. β-galactosidase activity was determined following the liquid culture protocol (Reynolds et al., 2001). All measurements were repeated at least three times.

## Results

### Identification of *MCH4* as a multicopy suppressor of glycine sensitivity

The parent strain for these experiments is of S288c background and is prototorophic for amino acids but requires uracil for growth (yJF176). Following a published protocol (Becker, D. M. and Lundblad, Victoria, 1993), the 1,580 basepairs upstream of the *GAP1* open reading frame were replaced with the kanMX selection marker followed by *P*_ADH1_; in addition, the first two lysines of Gap1p, 9 and 16, were mutated to arginines (the deletion of the extended *GAP1* promoter is essential as it prevents rejection of *GAP1* by homologous recombination that occurs at high frequency (Gresham et al., 2010)). In agreement with a previous report (Risinger et al., 2006), the resulting strain (yAM92) was sensitive to submillimolar concentrations of several amino acids when grown on synthetic medium (AmmSC). In particular, 200 μM glycine was sufficient to completely block growth on plates. This strain was transformed with a library of genomic DNA inserts on a multicopy plasmid (*URA3* selection marker, 5-10 kb inserts of genomic DNA from wild type yeast (FY4) constructed according to (Liu, 2002)), and resistant clones were selected on AmmSC + 200 μM glycine. Out of approximately 40,000 transformants, 10 glycine resistant clones were identified. Of these, 5 required plasmid for growth and were found to contain fragments of chromosome XV between positions 89,000 and 99,000; subcloning led to the conclusion that a piece of chromosome XV between positions 94,718 and 99,118 containing genes YOL119C (*MCH4*) and YOL118C (a dubious open reading frame) was sufficient to confer glycine resistance.

To confirm the role of *MCH4* in resistance to glycine, we constructed strains that either lacked Mch4p or overexpressed this protein in the background of *GAP1*^*UR*^ and compared them in plate growth assays. As shown in Figure 1A, Mch4p overexpression strain demonstrates increased resistance to glycine while *mch4Δ* strain is hypersensitive to this amino acid. This result also demonstrated that YOL118C is not involved in glycine resistance since in the overexpression strain *P*_ADH1_ replaced YOL118C while this open reading frame is still intact in *mch4Δ*. The protective effect of Mch4p is specific to glycine since overexpression or deletion had no effect on growth in the presence of lysine (Figure1B) or any other amino acid tested (Figure S1). Investigation of growth in liquid batch cultures showed that overexpression of Mch4p has an immediate protective effect: while addition of 500 μM glycine completely arrests growth within 30 minutes, *P*_ADH1_-Mch4 cells continue growing without a lag phase albeit at a slightly slower rate (Figure S2).

### Regulation of MCH4

No function has been previously assigned to *MCH4*. This nonessential gene codes for a protein homologous to mammalian monocarboxylate transporters (Lafuente and Gancedo, 1999); however, *MCH4* is not activated by growth on lactate, pyruvate, acetate, or any other carbon source tested, and no uptake or secretion of monocarboxylates could be detected in a *mch4Δ* strain (Makuc et al., 2001). According to the transcription factor occupancy data (MacIsaac et al., 2006), *MCH4* promoter binds Gcn4 and Gln3, transcription factors implicated in nitrogen catabolite repression. However, microarray analysis did not identify *MCH4* as a subject of nitrogen catabolite repression (Godard et al., 2007). Other transcription factors that bind to *P*_MCH4_ are Cad1p and Yap1p (stress response), Yap7p (unknown function) and Pho2p (an activator involved in several metabolic pathways). Previous reports indicated that *MCH4* is upregulated in response to osmotic and possibly other kinds of stress (Causton et al., 2001; Jelinsky et al., 2000; Epstein et al., 2001; Vanacloig-Pedros et al., 2016).

To study regulation of *MCH4* by the quality of nitrogen source as well as stress factors, we constructed a fusion of *P*_MCH4_ (including the first 48 amino acids of Mch4p) to β-galactosidase from *E. coli* on a low copy plasmid. The results of β-galactosidase activity measurements presented in Figure 2 show that *MCH4* is weakly regulated by the quality of nitrogen source as expression steadily increases in the row ammonia < allantoin < proline < glycine. We were surprised to find that glycine had a modest effect on expression of *MCH4* either in wild type or in an amino acid sensitive strain. While this finding is unexpected in view of our glycine toxicity suppression data, it is consistent with microarray results obtained in a study of glycine catabolism regulon (Gelling et al., 2004). The fact that *MCH4* is not regulated by glycine suggests that the primary function of this protein does not include transport of glycine. We also tested induction of *MCH4* by several stress factors (oxidation, heat, and high osmolarity and salinity) and found that *MCH4* was induced by osmotic stress (NaCl or sorbitol) but not by heat or oxidative stress (H_2_O_2_).

### Possible connection between response to amino acid toxicity and osmotic shock

The observation that *MCH4* is upregulated in high osmolarity medium suggested that there may be similarities between cellular response to osmotic shock and to high influx of amino acids. Therefore, we tested involvement of *MCH4* in osmotic shock response but found that overexpression or deletion of this gene had no effect on growth in the presence of 1 M NaCl or 1.5 M sorbitol (data not shown). However, a strain with disrupted HOG signaling pathway (yAM119; *pbs2Δ*) which is known to be hypersensitive to high osmolarity (Hohmann, 2002) turned out to be slightly more resistant to lysine but not glycine (Figure S3). While this effect was reproducible, it was not strong enough to draw any definitive conclusions.

### Contribution of *AQR1, ATR1, QDR1, QDR2*, and *QDR3* to amino acid resistance

It has been suggested that several yeast transporters from the major facilitator superfamily (MFS) are able to export amino acids from the cell. In particular, *AQR1, ATR1, QDR1, QDR2*, and *QDR3* have been proposed for this role (Sa-Correia et al., 2009) based on the following observations: i) they are subject to nitrogen catabolite repression, just like many other genes involved in amino acid metabolism; ii) they are coregulated with known exporters of polyamines and other toxic compounds; iii) one of the genes from this family is the only known amino acid exporter (*AQR1* is associated with threonine export (Velasco et al., 2004)). Logically, an exporter of amino acids should contribute to resistance to amino acids in the *GAP1*^*UR*^ background. Therefore, we constructed strains with deletions of *AQR1, ATR1, QDR3*, and *QDR2*-*QDR1* and tested their ability to grow in the presence of toxic amount of glycine, lysine, histidine, isoleucine, and threonine but detected at best very small changes in sensitivity to any of the tested amino acids (Figure S4).

### Amino acid catabolism genes suppress toxicity of branched chain amino acids

We have also attempted to clone multicopy suppressors of amino acid toxicity for lysine, cysteine, histidine, and isoleucine following the procedure described above for glycine. In each case, 40,000 transformants were screened but this procedure failed to identify any genes conferring resistance to lysine, cysteine, or histidine. In the case of isoleucine, we identified 22 clones of which 18 required plasmid for growth. These clones were found to contain either *LEU4* (6 clones) or *BAT2* (12 clones) inserts; subcloning confirmed that fragments containing only *LEU4* or *BAT2* were sufficient to confer resistance to 2 mM isoleucine. *BAT2* codes for a cytosolic branched chain amino acid aminotransferase, an enzyme that catalyzes the first step in catabolism of branched chain amino acids. *LEU4*, an alpha-isopropylmalate synthase, catalyzes the first step in leucine biosynthesis. Based on these findings, we conclude that our approach most readily identified genes involved in catabolism of the toxic amino acid (isoleucine) and failed to produce results for amino acids that cannot be catabolised.

## Discussion

To conclude, amino acid toxicity in yeast is an interesting and understudied phenomenon. While it is most pronounced under synthetic conditions (overexpression of unregulated Gap1p), it is likely present in natural context as evidenced by numerous reports of amino acid secretion under conditions of perturbed metabolism (Casalone et al., 1997; Feller et al., 1999; Hess et al., 2006; Lewis and Phaff, 1964; Delgado et al., 1982; Velasco et al., 2005), peptide uptake (Melnykov, 2016), or simply upon entry into the stationary phase of growth (Paczia et al., 2012). Among several phenotypes sensitive to amino acids (Ljungdahl et al., 1992; Risinger et al., 2006; Ruiz et al., 2017), we chose the *GAP1*^*UR*^ strain for our genetic selection experiment as the one with a broad spectrum of sensitivity.

Selection of multicopy suppressors of glycine toxicity identified *MCH4*. Based on the experiments described above, we cannot assign a function to this gene but our data supports the idea that it is not involved in glycine transport. The subcellular localization of Mch4p remains unknown. It was previously reported that Mch4p tagged with GFP at the C-terminus localized to the vacuole (Makuc et al., 2001) although no functional characterization of the fusion protein was carried out. We constructed a similar fusion and determined that it was not functional in glycine toxicity spotting assay. Interestingly, another transporter from the MCH family (*MCH3* also known as *ESBP6*) was recently identified in a screen for multicopy suppression of lactic acid toxicity even though *MCH3* was not regulated by exposure to lactic acid (Sugiyama et al., 2016). Thus, at least two MCH transporters of budding yeast contribute to stress resistance while their physiological function is still unknown. More generally, it has been deduced from large scale studies of yeast resistance to various drugs that many transporters confer resistance to a compound that does not resemble their natural substrates (Venancio et al., 2010). The only *S. cerevisiae* MCH family protein with a defined role (*MCH5*) serves as a transporter of riboflavin (Reihl and Stolz, 2005).

The mechanism of amino acid toxicity in the *GAP1*^*UR*^ strain is still not understood. Several mechanisms such as dissipation of the proton gradient, induction of the GCN pathway, and mischarging of tRNAs were considered in previous reports (Risinger et al., 2006; Ruiz et al., 2017). Based on our result in the mutant with disrupted HOG signaling pathway, we can add osmotic shock to this list. Yet another possible mechanism is formation of amyloid-like fibrils which has been shown to take place for glycine in vitro (Banik et al., 2016) and for other amino acids both *in vitro* and *in vivo* (Shaham-Niv et al., 2015; Laor et al., 2019). It seems likely though that different amino acids are toxic for different reasons. The mechanisms that are used by yeast to cope with unlimited amino acid uptake are also not clear. Certainly, for those amino acids that can be efficiently catabolized (Homann et al., 2005), degradation is the primary coping mechanism. In addition, considering multiple reports of amino acid excretion, it seems likely that yeast utilize this mechanism as well. Despite this, we did not find any evidence for involvement of several putative amino acid exporters in dealing with amino acid stress in the *GAP1*^*UR*^ strain. Future work aimed at identifying natural substrates of Mch4p will clarify the physiological function of this transporter and might shed light on mechanisms of glycine toxicity.

## Acknowledgments

Funding was provided by National Institutes of Health (Grant No. 1 R01 HL109505).

---

**Figure S1.**
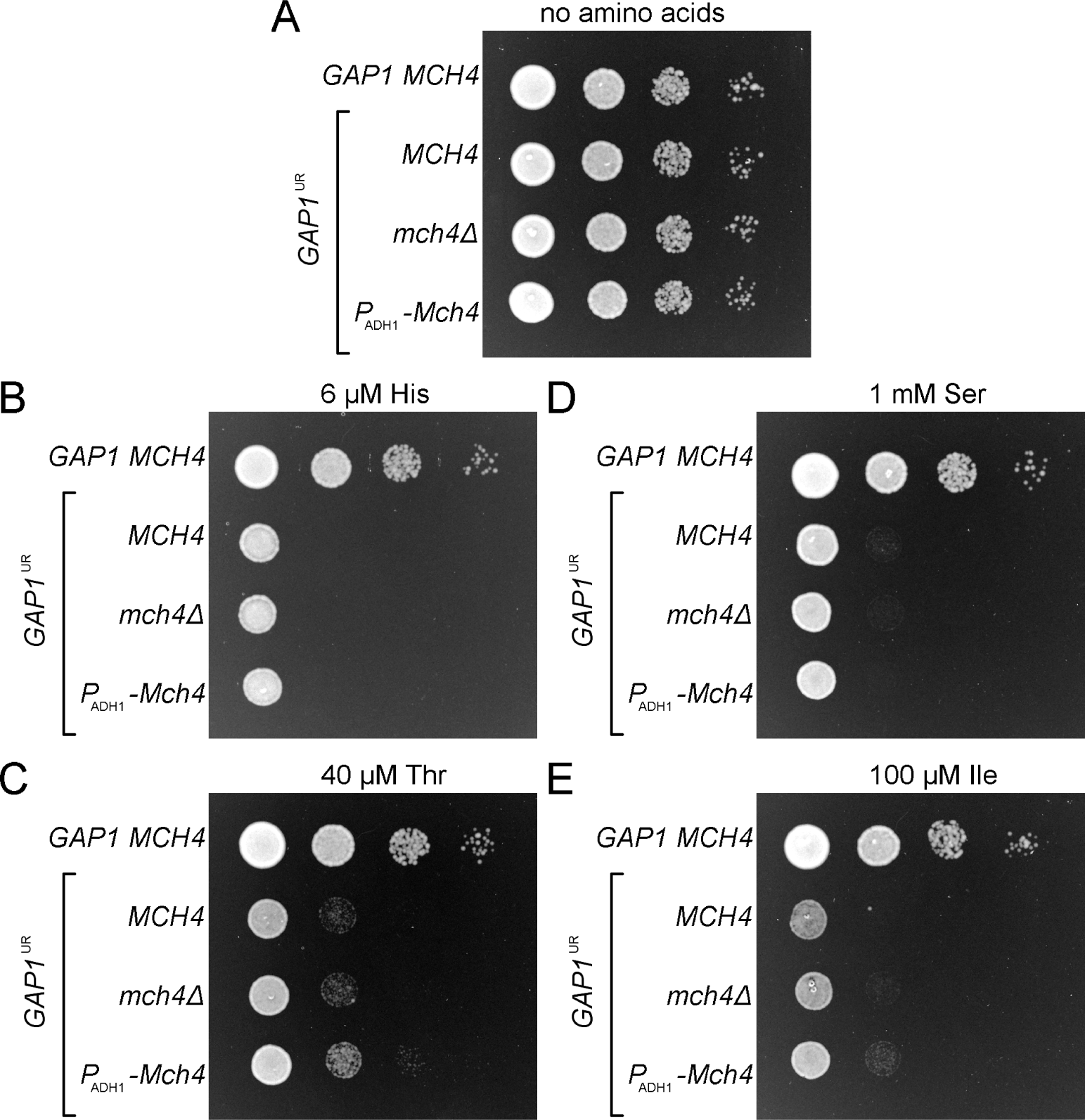
Toxicity of low concentrations of amino acids (A, no amino acids; B, 6 μM histidine; C, 40 μM threonine; D, 1 mM serine; E, 100 μM isoleucine). For each panel, the rows of dilutions from top to bottom are the wild type strain (yJF176) followed by strains sensitive to amino acids and varying in *MCH4* locus: wild type *MCH4* (yA92), deleted (yA93), and overexpressed from the *ADH1* promoter (yA94). The cells were grown on complete synthetic medium (AmmSC) supplemented with indicated concentration of amino acid.

**Figure S2.**
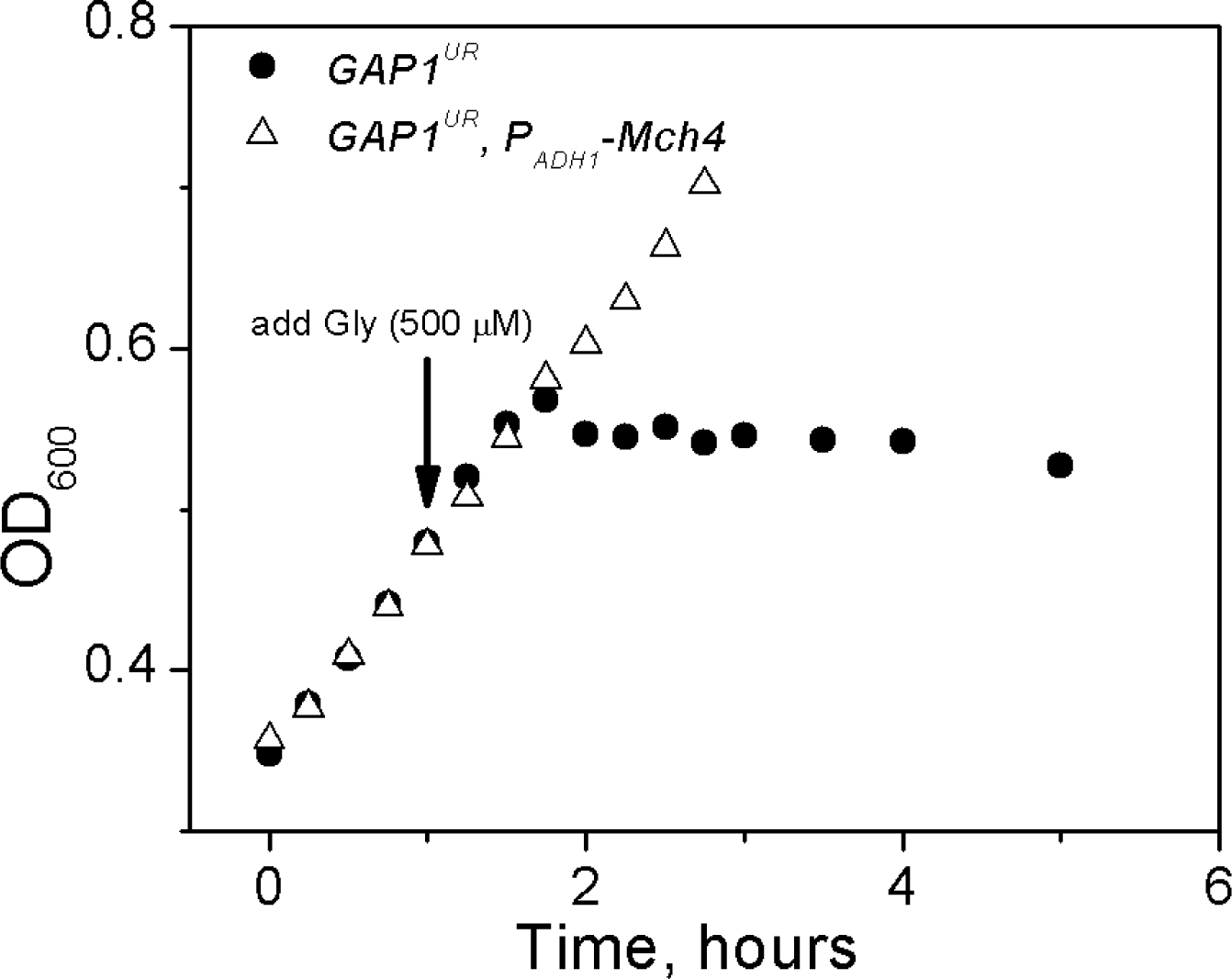
Effect of glycine addition (500 μM) on growth of yeast strain hypersensitive to amino acids with either wild type *MCH4* locus (yA92, •) or overexpressing Mch4p (yA94, Δ). The cells were grown on complete synthetic medium (AmmSC).

**Figure S3.**
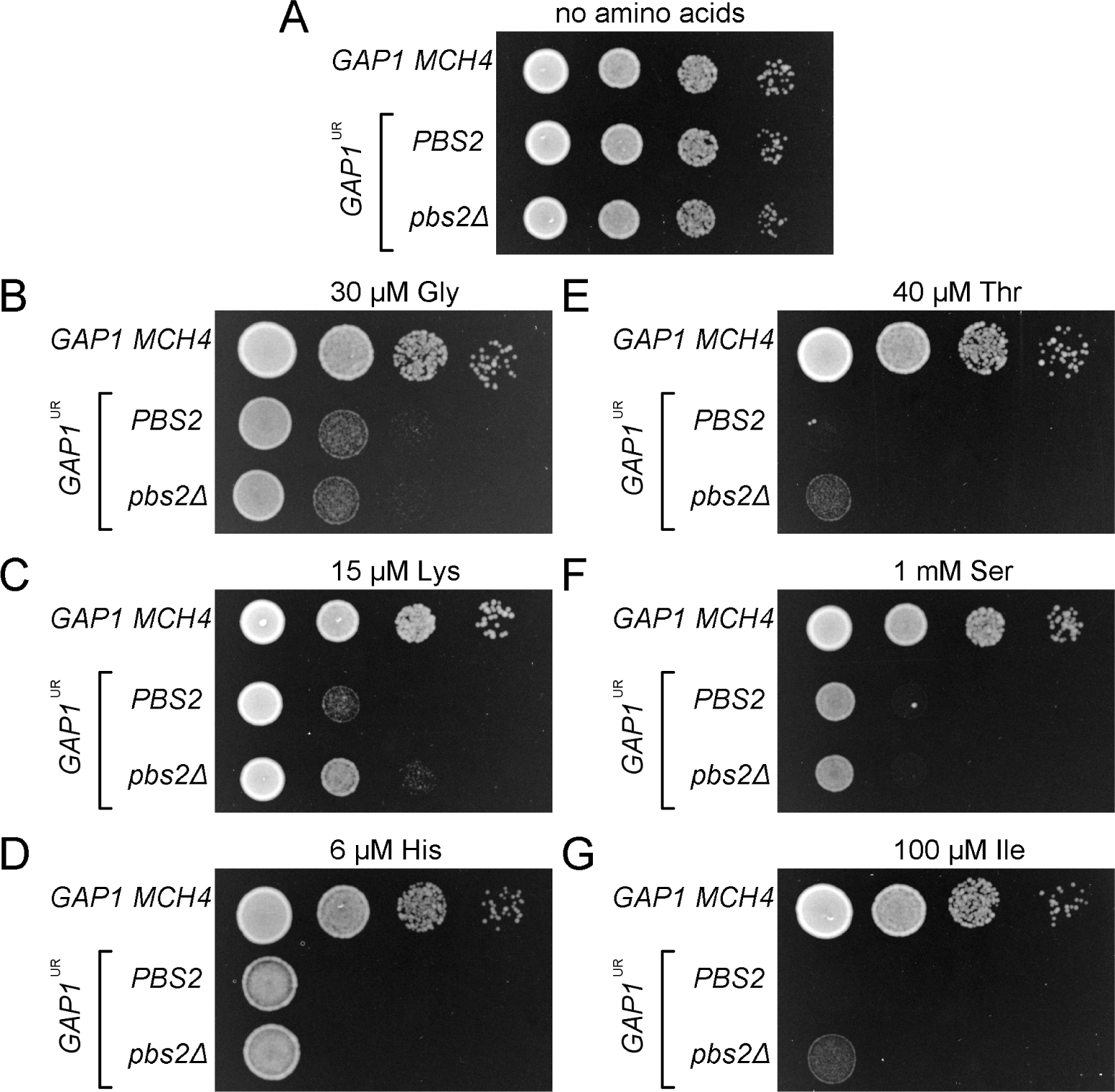
Effect of HOG pathway disruption on sensitivity to amino acids. (A, no amino acids; B, 30 μM glycine; C, 15 μM lysine; D, 6 μM histidine; E, 40 μM threonine; F, 1 mM serine; G, 100 μM isoleucine). For each panel, the rows of dilutions from top to bottom are the wild type strain (yJF176) followed by strains sensitive to amino acids with functional (yA92, *PBS2*) and disrupted (yA119, *pbs2*Δ) HOG pathway. The cells were grown on complete synthetic medium (AmmSC) supplemented with indicated concentration of amino acid.

**Figure S4.**
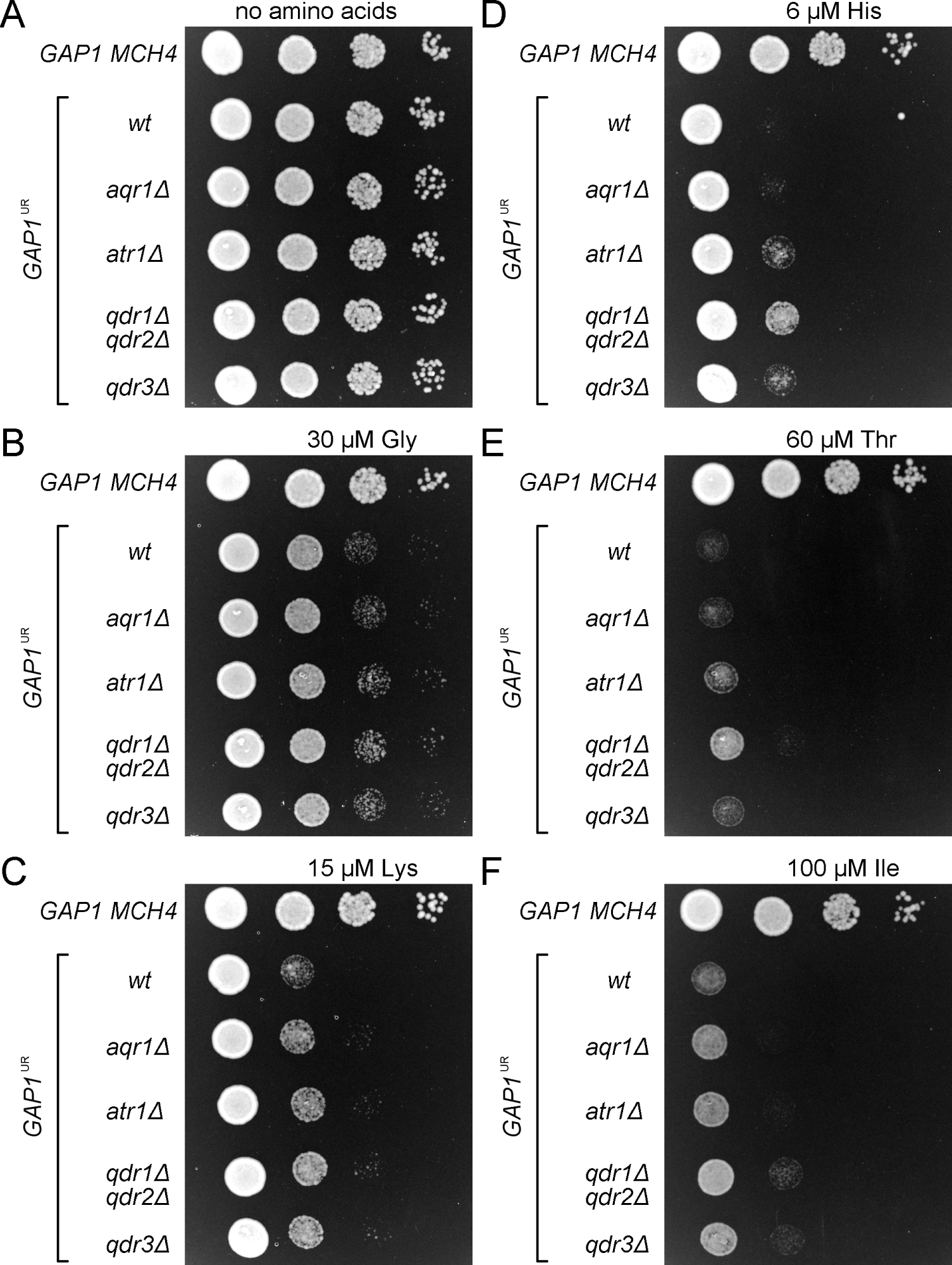
Effect of deletion of several MFS transporters on sensitivity to amino acids. (A, no amino acids; B, 30 μM glycine; C, 15 μM lysine; D, 6 μM histidine; E, 60 μM threonine; F, 100 μM isoleucine). For each panel, the rows of dilutions from top to bottom are the wild type strain (yJF176) followed by strains sensitive to amino acids with intact MFS transporters (yA92) or deleted *aqr1* (yA96), *atr1* (yA97), *qdr1* and *qdr2* (yA99), or *qdr3* (yA100). The cells were grown on complete synthetic medium (AmmSC) supplemented with indicated concentration of amino acid.

